# Limited Cross-Species Virus Transmission in a Spatially Restricted Coral Reef Fish Community

**DOI:** 10.1101/2022.05.17.492384

**Authors:** Vincenzo A. Costa, David R. Bellwood, Jonathon C.O. Mifsud, Jemma L. Geoghegan, Edward C. Holmes, Erin Harvey

**Affiliations:** Sydney Institute for Infectious Diseases, School of Life and Environmental Sciences and Sydney Medical School, The University of Sydney, Sydney, NSW 2006, Australia; Research Hub for Coral Reef Ecosystem Functions, College of Science and Engineering, James Cook University, Townsville, QLD 4811, Australia; Department of Microbiology and Immunology, University of Otago, Dunedin 9016, New Zealand; Institute of Environmental Science and Research, Wellington 5022, New Zealand

## Abstract

The Great Barrier Reef (GBR) – the largest coral reef ecosystem in the world – supports over 1200 fish species with some of the highest population densities and diversities seen in vertebrates, offering a high potential for virus transmission among species. As such, the GBR represents an exceptional natural ecosystem to determine the impact of host community diversity on virus evolution and emergence. In recent decades the GBR has also experienced significant threats of extinction, making it one of the most vulnerable ecosystems on the planet. However, our understanding of virus diversity and connectivity in tropical reef fishes remains poor. Here, we employed metatranscriptomic sequencing to reveal the viromes of 61 reef fish species. This identified a total of 132 viruses, 38 of which were vertebrate-associated and therefore likely infecting the fish, including a novel isolate of *Santee-cooper ranavirus* (*Iridoviridae*). Notably, we found little evidence for virus transmission between fish species living within a very restricted geographical space – a 100 m^2^ coral reef ecosystem – suggesting that there might be important host genetic barriers to successful cross-species transmission despite regular exposure. We also identified differences in virome composition between reef fish families, such that cryptobenthic reef fishes – characterized by small body sizes and short life-spans – exhibited greater virome richness compared to large reef fishes. This study suggests that there are important barriers to cross-species transmission, and that successful emergence in a reef fish community likely requires active host adaptation, even among closely related host species.

## Introduction

Despite their long evolutionary history, extensive diversity and complex ecological interactions, fish are severely under-sampled in studies of viral ecology and evolution. Economically, fish represent approximately 401 billion USD to the global economy and supply a yearly average of 20.5 kilograms per capita for consumption [1]. Yet fish face continual and potentially ruinous threats from emerging viral infections [2]. Climate-associated changes in species interactions are likely to have devastating ecological and economic consequences, particularly in the context of infectious disease emergence [3]. It was recently estimated that climate change may be responsible for the extinction of almost half of the economically important fish species in the tropical Pacific region by 2100 [4].

Tropical coral reefs are particularly vulnerable to biodiversity loss. In recent decades, global warming has caused numerous coral bleaching events worldwide, with detrimental cascading effects on reef ecosystem functioning and biodiversity [5, 6, 7]. While tropical coral reefs make up only a small fraction of the marine environment, they support enormous biodiversity, accounting for approximately one-third of all currently described marine fish species [8]. Many are ‘cryptobenthic reef fishes’, characterized by small body sizes (i.e. adult sizes of approximately five centimetres), short life-spans, cryptic behaviour and benthic positioning on coral reefs [9, 10]. These diverse reef fish assemblages are of significant economic and cultural value to humans through aquaculture, fisheries, tourism and the aquarium trade [10, 11, 12].

Despite their economic and socio-ecological importance, little is known about the natural diversity of viruses that infect reef fishes, and how ecological and phylogenetic variability within a reef fish community impacts cross-species virus transmission and disease emergence. Host community diversity likely plays a central role in virus emergence, as contact between donor and recipient hosts is a prerequisite for virus transmission [13]. Yet, revealing the exact nature of that role has proven challenging [14]. On one hand, a high diversity of hosts could provide more transmission opportunities for viruses, thereby elevating the risk of disease emergence. Conversely, increased host community diversity may reduce the probability of disease emergence through the ‘dilution effect’, in which species richness provides more possible hosts for pathogens, in turn reducing disease occurrence in some species [14, 15]. As tropical coral reefs are considered biodiversity hotspots, they serve as an ideal natural ecosystem for exploring the impact of host diversity on the extent, pattern and evolution of virus diversity. Indeed, reef fishes display some of the highest densities and highest diversities of potential vertebrate hosts on the planet. In our study location, a standard sampling area of 3.5 m^2^ consistently supports between 50 to 150 fishes belonging to 15 to 25 species [74]. Furthermore, this density and richness is sustained year-round [75], offering a continual potential for cross-species transmission and high individual densities all in a readily transmitting aquatic medium.

Much of our knowledge on the tropical reef virosphere is skewed towards those viruses associated with coral species and their symbionts [16]. While epizootic infections have previously been reported in tropical reef fishes across the Western Atlantic and Gulf of Mexico [17], there is minimal data on viral diversity in fish from the Great Barrier Reef (GBR), Australia, although this is the largest and among the most threatened reef ecosystems in the world [5, 6, 18]. A recent metatranscriptomic analysis of the pygmy goby (*Eviota zebrina*) identified three viruses (families *Arenaviridae, Hantaviridae, Paramyxoviridae*) despite *Eviota* exhibiting maximum lifespans of between 60 and 100 days (including a 24 to 26 day pelagic larval phase) [19, 20].

The rate of infectious disease emergence is expected to increase in marine environments, particularly in the context of climate change [3]. As ocean temperatures continue to rise, many Eastern Australian tropical fishes have begun to shift their distribution poleward to temperate reefs situated at higher latitudes, through tropicalization [21, 22]. While the broad-scale ecological impacts of tropicalization are becoming increasingly apparent in native temperate fishes [22], it is unclear how this will impact virus ecology and infectious disease emergence.

Tropical ornamental fishes may also act as viral vectors of disease in economically important farmed species. For example, in Australia, dwarf gourami (*Trichipodus lalius*) can transmit infectious kidney and spleen necrosis virus (genus *Megalocytivirus, Iridoviridae*) to domestic fishes, often with detrimental impacts including disease outbreaks in iconic species such as Murray cod (*Maccullochella peelii*) [23, 24]. Moreover, tropical wrasses (Labridae) have been considered effective biological control agents in aquaculture for their natural ability to consume pests [25]. However, temperate cleaner wrasses have been reported to be important drivers of outbreaks of viral haemorrhagic septicaemia virus (*Rhabdoviridae*) in farmed salmonids [26]. As such, revealing viral diversity in ornamental tropical reef fishes is imperative to understanding the risk of disease emergence in both wild and domestic fish populations.

Revealing the nature of viral transmission within and among species in natural environments is central to our understanding of disease emergence. With exceptionally high species diversity and highly variable individual abundances and ecologies, coral reefs offer an exciting opportunity to explore cross-species virus transmission in a complex natural high-diversity ecosystem. Our goal, therefore, was to employ total RNA-sequencing (i.e. metatranscriptomics) to reveal the diversity, abundance and composition of viruses infecting reef fishes from the GBR, utilising a fish community from a 100 m^2^ coral reef ecosystem. In particular, we aimed to: (i) reveal viral diversity and evolution in tropical reef fishes, (ii) identify how often viruses are exchanged in a spatially restricted reef fish community, (iii) determine whether there are differences in virome composition between reef fish families, as well as between cryptobenthic reef fishes and large reef fishes, that differ enormously in size, metabolic rate, lifespan and fecundity [10], and (iv) identify novel viruses that may pose an emerging threat to Australian fisheries, aquaculture and the aquarium trade.

## Materials and Methods

### Animal ethics

Fish were collected under a Great Barrier Reef Marine Park Authority permit (G16/37684.1) and James Cook University Animal Ethics permit A2752.

### Tropical reef fish sample collection

Seemingly healthy fishes (n = 193) were collected in early April 2021 at Orpheus Island, GBR (18°36’44.3”S 146°28’59.4”E). These included 61 species across 16 reef fish families: Gobiidae (gobies) (n = 30 species), Labridae (wrasses) (n = 6), Pomacentridae (damselfishes) (n = 5), Blenniidae (blennies) (n = 5), Acanthuridae (surgeonfishes) (n = 3), Apogonidae (cardinalfishes) (n = 2), Monacanthidae (filefishes) (n = 2), Tetraodontidae (pufferfishes) (n = 1), Pseudochromidae (dottybacks) (n = 1), Chaetodontidae (butterflyfishes) (n = 1), Atherinidae (silversides, hardyheads) (n = 1), Serranidae (groupers) (n = 1), Tripterygiidae (triplefin blennies) (n = 1), Muraenidae (moray eels) (n = 1), Bythitidae (brotulas) (n = 1) and Ophichthidae (snake eels) (n = 1). (Supplementary Table 1). Importantly, of these species, 42 were collected from the reef fish community within a 100 m^2^ sampling area (along the northern margin of Pioneer Bay) (Supplementary Table 1). All fish caught were euthanized using clove oil, transported to the lab on ice, then placed either dissected (liver and gills) or whole in RNAlater. Specimens were then stored at −80°C until RNA extraction.

**Table 1.** Description of viruses identified in this study.

| Virus name | No.<br>of<br>reads | Virus family | Host | No. of<br>ORFs/Segments | Length (nt) | Complete<br>genome |
| --- | --- | --- | --- | --- | --- | --- |
| Goby astrovirus 2 | 24100 | *Astroviridae* | *Cabillus tongarevae* | 3 | 7226 | Y |
| Goby astrovirus 2 | 2767 | *Astroviridae* | *Istigobius decoratus* | 3 | 7085 | Y |
| Goby astrovirus 2 | 9023 | *Astroviridae* | *Istigobius nigroocellatus* | 3 | 7155 | Y |
| Goby astrovirus 2 | 1184 | *Astroviridae* | *Asterropteryx*<br>*semipunctatus* | 3 | 7045 | Y |
| Goby astrovirus 1 | 2863 | *Astroviridae* | *Istigobius goldmanni* | 3 | 7087 | Y |
| Goby astrovirus 3 | 6709 | *Astroviridae* | *Luposicya lupus* | 2 | 7355 | Y |
| *Blenniella* astrovirus | 59 | *Astroviridae* | *Blenniella* sp. | 1 | 744 | N |
| *Salarius guttatus* piscichuvirus | 40334 | *Chuviridae* | *Salarias guttatus* | 3 | 10801 | Y |
| *Abudefduf bengalensis* circovirus | 54 | *Circoviridae* | *Abudefduf bengalensis* | 1 | 873 | N |
| *Enneapterygius tutuilae* letovirus | 26 | *Coronaviridae* | *Enneapterygius tutuilae* | 1 | 856 | N |
| *Chaetodon aureofaciatus* hepacivirus | 1029 | *Flaviviridae* | *Chaetodon aureofaciatus* | 1 | 8826 | Y |
| *Luposicya lupus* actinovirus | 6144 | *Hantaviridae* | *Luposicya lupus* | 3 | 11533 | Y |
| Pygmy goby hantavirus | 972 | *Hantaviridae* | *Eviota zebrina* | 3 | 5894 | N |
| *Trimma fangi* actinovirus | 106 | *Hantaviridae* | *Trimma fangi* | 2 | 1319 | N |
| *Pleurosicya* actinovirus | 123 | *Hantaviridae* | *Pleurosicya* sp. | 2 | 2103 | N |
| *Cephalopholis boenak* actinovirus | 9 | *Hantaviridae* | *Cephalopholis boenak* | 1 | 1029 | N |
| *Istigobius decoratus* hepevirus | 16 | *Hepeviridae* | *Istigobius decoratus* | 1 | 393 | N |
| *Halichoeres melanurus* ranavirus | 9835 | *Iridoviridae* | *Halichoeres melanurus* | 9 | 10821 | N |
| *Enneapterygius tutuilae* iridovirus | 35545 | *Iridoviridae* | *Enneapterygius tutuilae* | 9 | 20970 | N |
| *Cheilodipterus quinquelineatus* orthomyxovirus | 979 | Orthomyxoviridae | *Cheilodipterus quinquelineatus* | 5 | 9090 | N |
| Pygmy goby paramyxovirus | 238 | Paramyxoviridae | *Eviota zebrina* | 1 | 1314 | N |
| *Enneapterygius tutuilae* paramyxovirus | 322 | Paramyxoviridae | *Enneapterygius tutuilae* | 1 | 618 | N |
| *Istigobius ornatus* parvovirus | 1746 | Parvoviridae | *Istigobius ornatus* | 3 | 4284 | Y |
| *Istigobius rigillus* parvovirus | 266 | Parvoviridae | *Istigobius rigillus* | 1 | 1656 | N |
| *Ecsenius stictus* parvovirus | 2707 | Parvoviridae | *Ecsenius stictus* | 2 | 4286 | Y |
| *Dinematichthys* parvovirus | 122 | Parvoviridae | *Dinematichthys* sp. | 1 | 1236 | N |
| *Luposicya lupus* ichthamaparvovirus | 12328 | Parvoviridae | *Luposicya lupus* | 1 | 1299 | N |
| *Blenniella* picornavirus | 2 | Picornaviridae | *Blenniella* sp. | 1 | 347 | N |
| *Enneapterygius tutuilae* picornavirus | 225 | Picornaviridae | *Enneapterygius tutuilae* | 1 | 3840 | N |
| *Exyrias* picornavirus | 44 | Picornaviridae | *Exyrias* sp. | 1 | 1373 | N |
| *Halichoeres marginatus* picornavirus | 27506 | Picornaviridae | *Halichoeres marginatus* | 1 | 8344 | Y |
| *Neopomacentrus bankieri* picornavirus | 2955 | Picornaviridae | *Neopomacentrus bankieri* | 1 | 8408 | Y |
| *Asterropteryx spinosa* picornavirus | 8307 | Picornaviridae | *Asterropteryx spinosa* | 1 | 8405 | Y |
| *Asterropteryx semipunctatus* picornavirus | 210 | Picornaviridae | *Asterropteryx semipunctatus* | 1 | 992 | N |
| *Pomacentrus moluccensis* picornavirus | 12 | Picornaviridae | *Pomacentrus moluccensis* | 1 | 321 | N |
| *Trimma okinawae* rhabdovirus | 13 | Rhabdoviridae | *Trimma okinawae* | 2 | 945 | N |
| *Trimma striatum* rhabdovirus | 2 | Rhabdoviridae | *Trimma striatum* | 1 | 488 | N |
| *Istigobius nigroocellatus* reovirus | 282 | Reoviridae | *Istigobius nigroocellatus* | 6 | 6966 | N |

### RNA extraction, library preparation and metagenomic next-generation sequencing

Tissue specimens were combined based on species (e.g. liver and gills) and total RNA was extracted using the RNeasy Plus Mini Kit (Qiagen, Hilden, Germany) as previously described in [18, 26]. RNA was quantified using a UV-Vis cuvette spectrophotometer (DeNovix, Delaware, United States) and a parallel capillary electrophoresis instrument (Fragment Analyzer) (Agilent, California, United States). RNA from each individual fish was then pooled within each species (Supplementary Table 1), resulting in a total of 61 RNA sequencing libraries. These libraries were constructed using the Truseq Total RNA Library Preparation Protocol (Illumina). Host ribosomal RNA (rRNA) was depleted with the Ribo-Zero Plus Kit (Illumina) and paired-end sequencing (150 bp) was carried out on the NovaSeq 500 platform (Illlumina). To reduce the impact of index hopping and false virus-host assignments, each library was sequenced on two different lanes. Library construction and metatranscriptomic sequencing were performed by the Australian Genome Research Facility (AGRF).

### Virus discovery pipeline and genome annotation

Illumina sequencing reads were first quality trimmed using Trimmomatic v.0.38 [28] then *de novo* assembled into contigs using MEGAHIT v.1.2.9 [29]. The resultant contigs for each library were used as a query against the NCBI nucleotide (nt) and non-redundant protein (nr) databases using BLASTn and Diamond (BLASTX) with an e-value search threshold of 1 × 10^−5^ [30]. Contigs with positive matches to viral sequences were inspected and predicted into open reading frames (ORFs) using Geneious Prime (v.2022.0) [31] (www.geneious.com). We first assigned a virus to a fish library if it was identified on both sequencing lanes. Next, to distinguish between those viruses infecting fish or those associated with diet or gill contamination (e.g. invertebrates, fungi, algae), we used our predicted viral sequences as a single query against the NCBI nt and nr databases using BLAST implemented in Geneious Prime. Using these results, we then assigned a virus as ‘vertebrate-associated’ based on sequence similarity to established vertebrate viruses on the NCBI databases, as well as e-value. Final confirmation was based on phylogenetic analysis (see below) as vertebrate and non-vertebrate viruses often exhibit a large degree of genetic divergence and hence are phylogenetically distinct [47, 51]. Viruses were defined as novel based on the broad criteria by The International Committee on Taxonomy of Viruses (https://talk.ictvonline.org). Viral genomes were annotated using the NCBI conserved domain (CDD) search and ‘Live Annotate and Predict’ tool in Geneious Prime, using related sequences obtained from NCBI/GenBank with a gene similarity cut-off of 25%.

### Phylogenetic analysis

To infer the evolutionary history of the viruses discovered here, we estimated family level phylogenetic trees using amino acid sequences of conserved genomic regions. These included RNA-dependent RNA polymerase (RdRp) for RNA viruses, and the DNA polymerase and major capsid protein for DNA viruses. We then aligned our novel viral sequences with those available on the NCBI/GenBank database (August 2021) using MAFFT v.7.450, employing the E-INS-i algorithm [32]. The amino acid sequence alignment was pruned using TrimAl v.1.2 to remove ambiguously aligned regions with a gap threshold of 0.9 and a variable conserve value [33]. The best-fit model of amino acid substitution was estimated with the ‘ModelFinder Plus’ (-m MFP) flag in IQ-TREE [34, 35]. Using these data, we then estimated phylogenetic trees using a maximum likelihood (ML) approach with 1,000 bootstrap replicates using IQ-TREE.

### Abundance estimation and virome statistical analysis

We employed RNA-seq by Expectation-Maximization (RSEM) (v1.2.28) to quantify the relative abundance of transcripts within each fish species transcriptome [36]. These included both viral genes and the stably expressed host reference gene, ribosomal protein S13 (RPS13). Abundance measures were standardised by dividing values against the total reads for each library. We calculated both alpha and beta diversity to compare virome composition between reef fish families, as well as between cryptobenthic reef fishes and large reef fishes. We also compared non-fish virome composition (i.e. those viruses associated with diet, environment or microbiome) as a form of internal control as these viruses are not impacted by aspects of host biology. Accordingly, we used Rhea scripts to calculate alpha diversity, including viral abundance, observed virome richness and Shannon diversity [37]. Statistical comparisons of alpha diversity were modelled using generalized linear models (GLMs) and tested using a likelihood-ratio test (χ^2^) and Tukey’s post hoc analysis with the *multcomp* package [38]. To compare viral communities between reef fish assemblages, we calculated beta diversity using a Bray–Curtis distance matrix with the *phyloseq* package [71]. These data were then tested using permutational multivariate analysis of variance (permanova) with the *vegan* package (adonis) [72]. All plots were constructed using *ggplot2* [39].

## Results

### Total diversity and abundance of viruses discovered in tropical reef fish

We sequenced 61 metatranscriptomes of tropical reef fish species for virus discovery, resulting in a median of 99,658,615 (range 45,583,181 – 121,905,422) reads and 371,944 (range 161,447 – 626,467) contigs per library. From this, we identified a total of 132 viruses infecting apparently healthy fish and their associated invertebrate and microbial species. This included 38 vertebrate-associated viruses, comprising 11 negative-sense single-stranded RNA viruses (−ssRNA) (*Hantaviridae, Chuviridae, Orthomyxoviridae, Rhabdoviridae, Paramyxoviridae*), 18 positive-sense single-stranded RNA viruses (+ssRNA) (*Astroviridae, Hepeviridae, Picornaviridae, Flaviviridae, Coronaviridae*), one double-stranded RNA virus (dsRNA) (*Reoviridae*), six single-stranded DNA viruses (ssDNA) (*Parvoviridae, Circoviridae*), and two double-stranded DNA viruses (dsDNA) (*Iridoviridae*) (Figure 1). Our metatranscriptomic analysis also identified 94 viruses that shared sequence similarity and phylogenetically clustered with viruses associated with invertebrate (arthropods, crustaceans, molluscs, platyhelminths, myriapods, nematodes) and fungal hosts (Supplementary Figures 1-4). These viruses were classified within the *Flaviviridae* (27% of total non-vertebrate virome abundance), *Narnaviridae* (24%), *Picornaviridae* (20%), *Nodaviridae* (10%), *Hepeviridae* (7%), *Solemoviridae* (4%), *Totiviridae* (1%), *Partitiviridae, Iflaviridae, Picobirnaviridae, Phenuiviridae, Qinviridae, Rhabdoviridae, Tombusviridae, Mimiviridae, Natareviridae, Jingchuvirales* (all less than 1%) [40, 41]. We also identified viruses that were related to negeviruses (2% of non-vertebrate virus abundance) [42] and quenyaviruses (<1%) [43] (Supplementary Figure 1). While many of these families include viruses that are known to infect fishes (e.g. *Flaviviridae, Picornaviridae, Nodaviridae, Hepeviridae, Totiviridae, Rhabdoviridae*), the viruses identified here were highly divergent and clustered with those that infect a broad range of invertebrate and fungal species (Supplementary Figures 1-4). We therefore focused our analysis on vertebrate-associated viruses.

**Figure 1.**
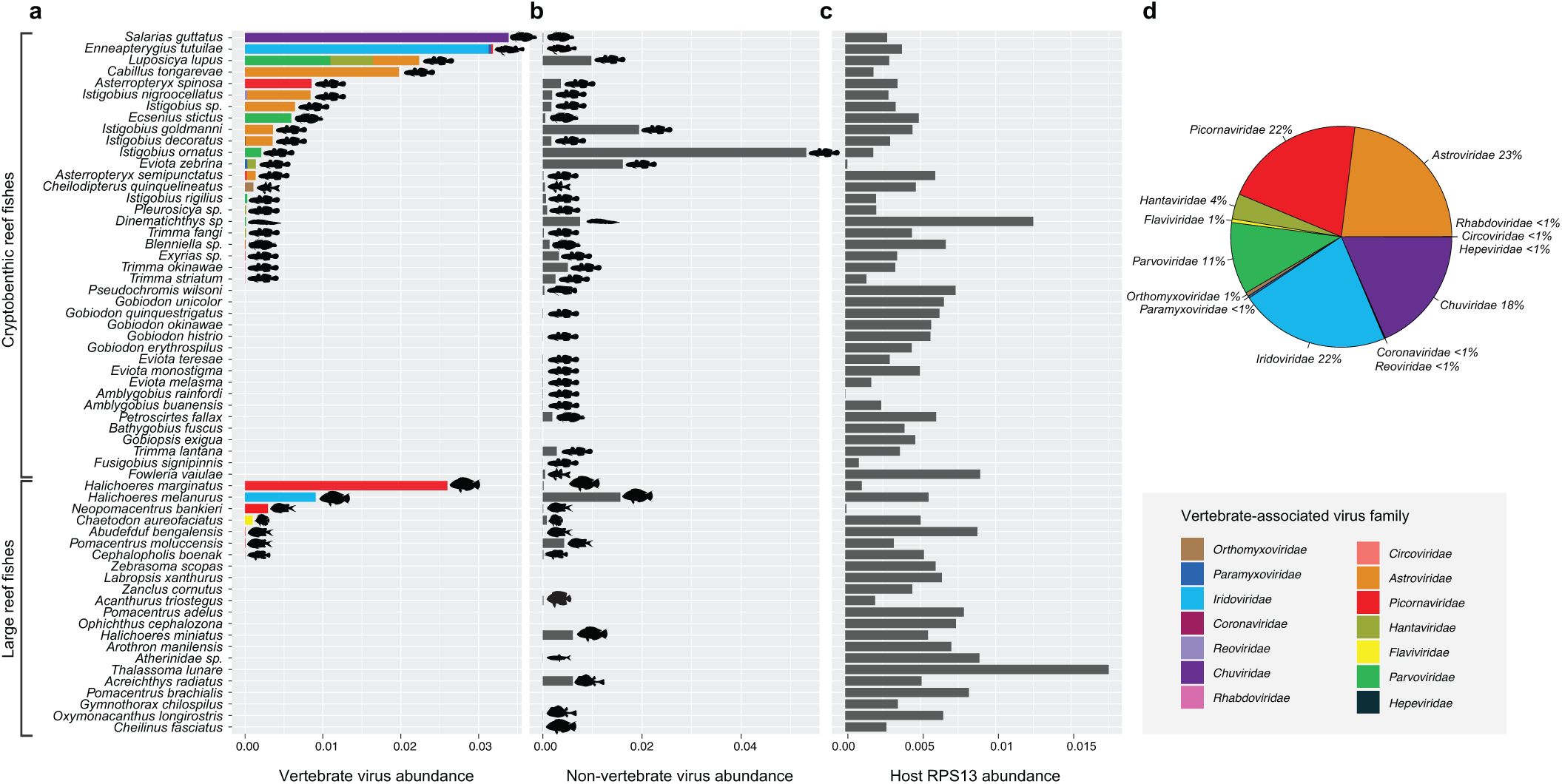
Standardized abundance of viral and host reads across reef fish libraries. Abundance (i.e., number of reads) of: (a) likely vertebrate-associated viruses; (b) non-vertebrate associated viruses (i.e., those from algae, fungi, coral, arthropods, crustaceans and protists); (c) host reference gene RPS13. Silhouettes represent the fish species and virus families are indicated by colour. (d) Total standardized abundance of vertebrate-associated viral families.

### Genome organisation and phylogenetic relationships of vertebrate-associated viruses *−ssRNA viruses*

Five actinoviruses were identified (subfamily *Actantavirinae, Hantaviridae*) (Figure 2). All of these were novel, with the exception for pygmy goby hantavirus, previously identified in *E. zebrina* [19]. We identified complete and partial actinovirus genomes in *Trimma fangi, Luposicya lupus, Cephalopholis boenak* and *Pleurosicya sp*. (Figure 2; Supplementary Table 2). The genome of *L. lupus* actinovirus contained all three expected actinovirus genomic regions (Figure 2). Notably, the S segment possessed an additional ORF upstream of the nucleoprotein in antisense orientation, also seen in the genome of perch actinovirus, but not in other actinoviruses [44].

**Figure 2.**
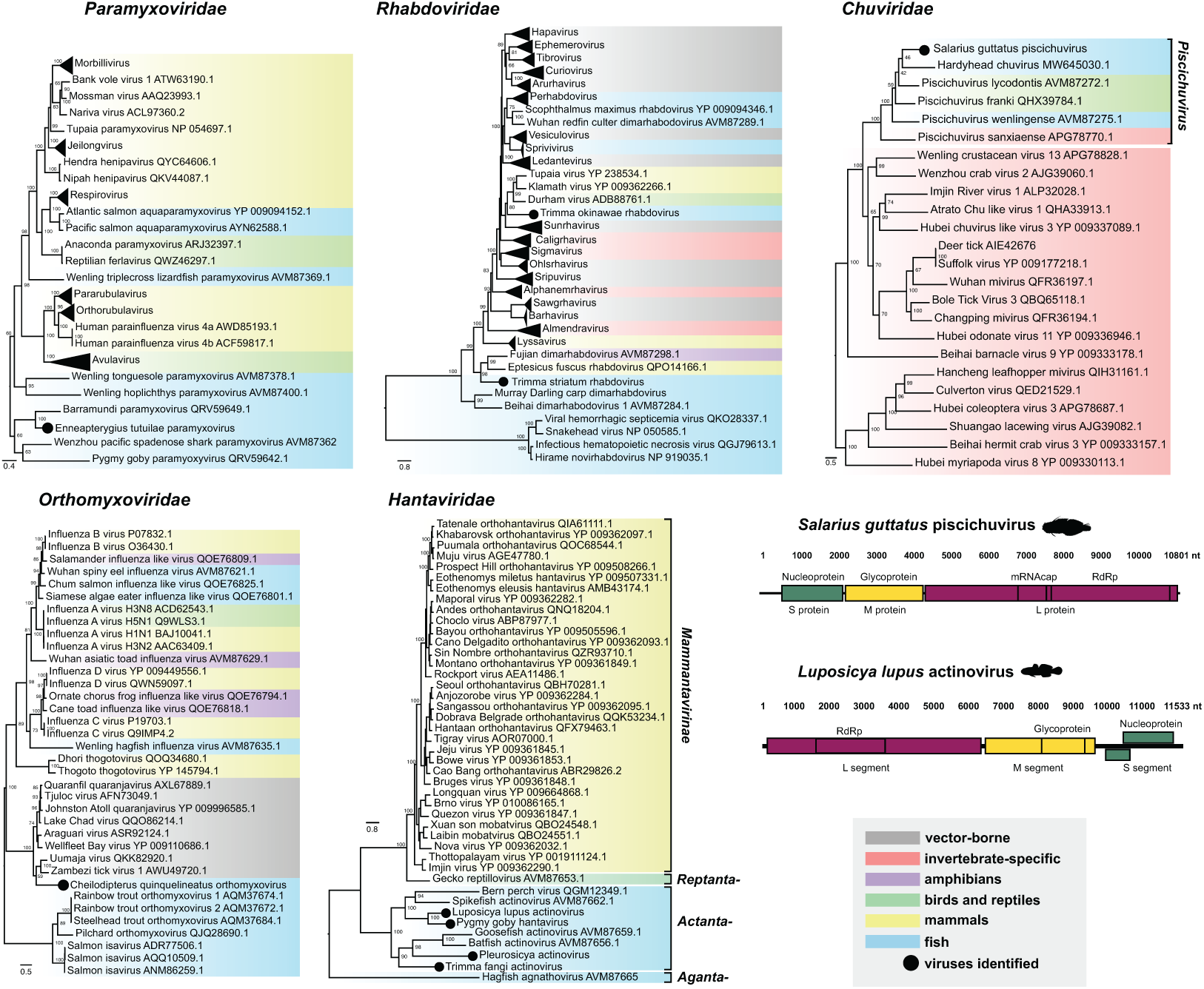
Genome structure and phylogenies of the RdRp gene of −ssRNA viruses identified in this study. Maximum likelihood phylogenies of novel and related virus species. Trees were midpoint rooted for clarity. Scale bar represents the number of amino acid substitutions per site. Tip labels represent virus name with NCBI/GenBank Accession. Tree branches are highlighted to broadly represent host taxonomy. Schematic genome diagrams illustrate genome orientation, length, predicted ORFs and gene products.

Two partial genomes of paramyxoviruses were identified, one in *E. zebrina* (i.e., pygmy goby paramyxovirus) [19] and the other in the *Enneapterygius tutuilae*. This virus was highly divergent from other classified ray-finned fish paramyxoviruses, including the genus *Aquaparamyxovirus* (Figure 2). We also identified partial genomes (L protein and nucleocapsid sequences) of two novel rhabdoviruses in *Trimma okinawae* and *Trimma striatum. T. okinawae* rhabdovirus fell within the monophyletic *Tupavirus* group that infects birds and mammals [45, 46], while *T. striatum* rhabdovirus clustered with the dimarhabdoviruses [47] (Figure 2). Among other −ssRNA viruses, we identified a novel pischichuvirus in *Salarius guttatus* and orthomyxovirus in *Cheilodipterus quinquelineatus* that showed sequence similarity to quaranjaviruses (Figure 2; Supplementary Table 2).

### +ssRNA viruses

Astroviruses were detected in seven fish species. All of these were members of the Gobiidae, except for *Blenniella sp*. (Blenniidae) which clustered with astroviruses that infect a broad range of ray-finned and jawless fish species (Figure 3). Similarly, we identified a divergent virus from this group in *L. lupus*, tentatively named goby astrovirus 3. This virus exhibited a genome of 7,355 nt with a predicted 5’ untranslated region (UTR) of 347 nt, two main ORFs, and a 3’ UTR of 347 nt. One ORF encoded the astrovirus ORF1a protein (protease) and the other, a polyprotein encoding RdRp and capsid protein. Notably, this arrangement is also seen in the genome of Wenling rattails astrovirus 5, but not in other currently described fish astroviruses that exhibit three main ORFs [47].

**Figure 3.**
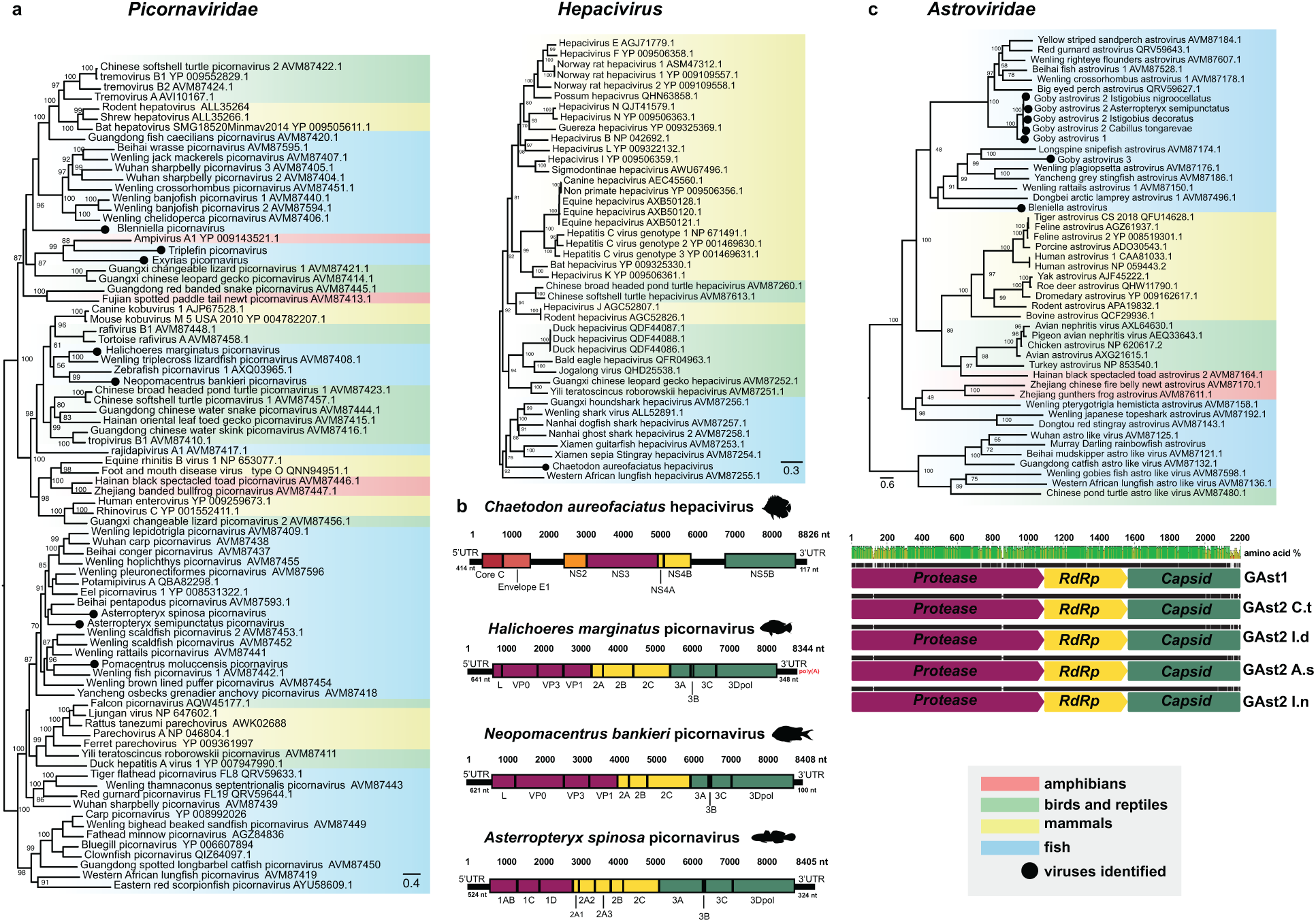
Genome organisation and phylogenies of the RdRp gene of +ssRNA viruses identified in this study. (a) Maximum likelihood phylogenies of the *Picornaviridae* and *Hepacivirus* (*Flaviviridae*), midpoint rooted for clarity. Scale bar represents the number of amino acid substitutions per site. Tip labels represent virus name with NCBI/Genbank Accession. Trees are highlighted to broadly illustrate host taxonomy. (b) Schematic genomes illustrate genome orientation, length and gene products. (c) Phylogeny of the *Astroviridae* and amino acid alignment of goby astrovirus 2 (GAst2) isolates against goby astrovirus 1 (GAst1). GAst2 are labelled to represent host species: GAst2 C.t = *C. tongaravae*, I.d = *I. decoratus*, A.s = *A. semipunctatus*, I.n = *I. nigroocellatus*.

The complete genomes of three novel picornaviruses and partial sequences of five novel picornaviruses were identified (Figure 3). Although these fish species were members of the same community, all eight viruses were highly divergent. Phylogenetic analysis of the 3D polymerase gene reveals clustering with other fish picornaviruses, except for *E. tutuilae* picornavirus and *Exyrias* picornavirus that clustered with *Ampivirus*, identified in the common newt (*Lissotriton vulgaris*) [48] (Figure 3).

We also identified the full genome of a novel hepacivirus (*Flaviviridae*) in *Chaetodon aureofaciatus* that contained conserved genomic regions – Core C, Envelope E1, NS2, NS3, NS4, NS5B – and fell within a distinct group of aquatic hepaciviruses [47] (Figure 3).

### dsRNA viruses

The near complete genome of a novel reovirus was identified in *Istigobius nigroocellatus*. This included six segments encoding the VP1 (guanyl transferase), VP2 (RdRp), VP3 (helicase), VP4 (NTPase), VP5 (outer capsid protein) and VP6 (core protein) proteins. Phylogenetic analysis of the VP2 gene revealed clustering within the *Aquareovirus* genus, as this virus shared 48-60% amino acid similarity with its closest relatives (Supplementary Table 2; Supplementary Figure 5).

### ssDNA viruses

We discovered a basal group of four parvoviruses that fell within the subfamily *Parvovirinae* (*Parvoviridae*). The genome of *Ecsensius stictus* parvovirus was a single contig of 4286 nt, containing two main ORFs. The left ORF was 1374 nt and encoded the conserved non-structural protein NS1 that has DNA helicase and ATPase function [65]. The right ORF encoded the structural VP1 protein (2571 bp). *I. ornatus* parvovirus exhibited a genome of 4284 nt and contained three main ORFs, one encoding NS1 (1878 nt) and the other two, encoding putative structural proteins (Figure 4).

**Figure 4.**
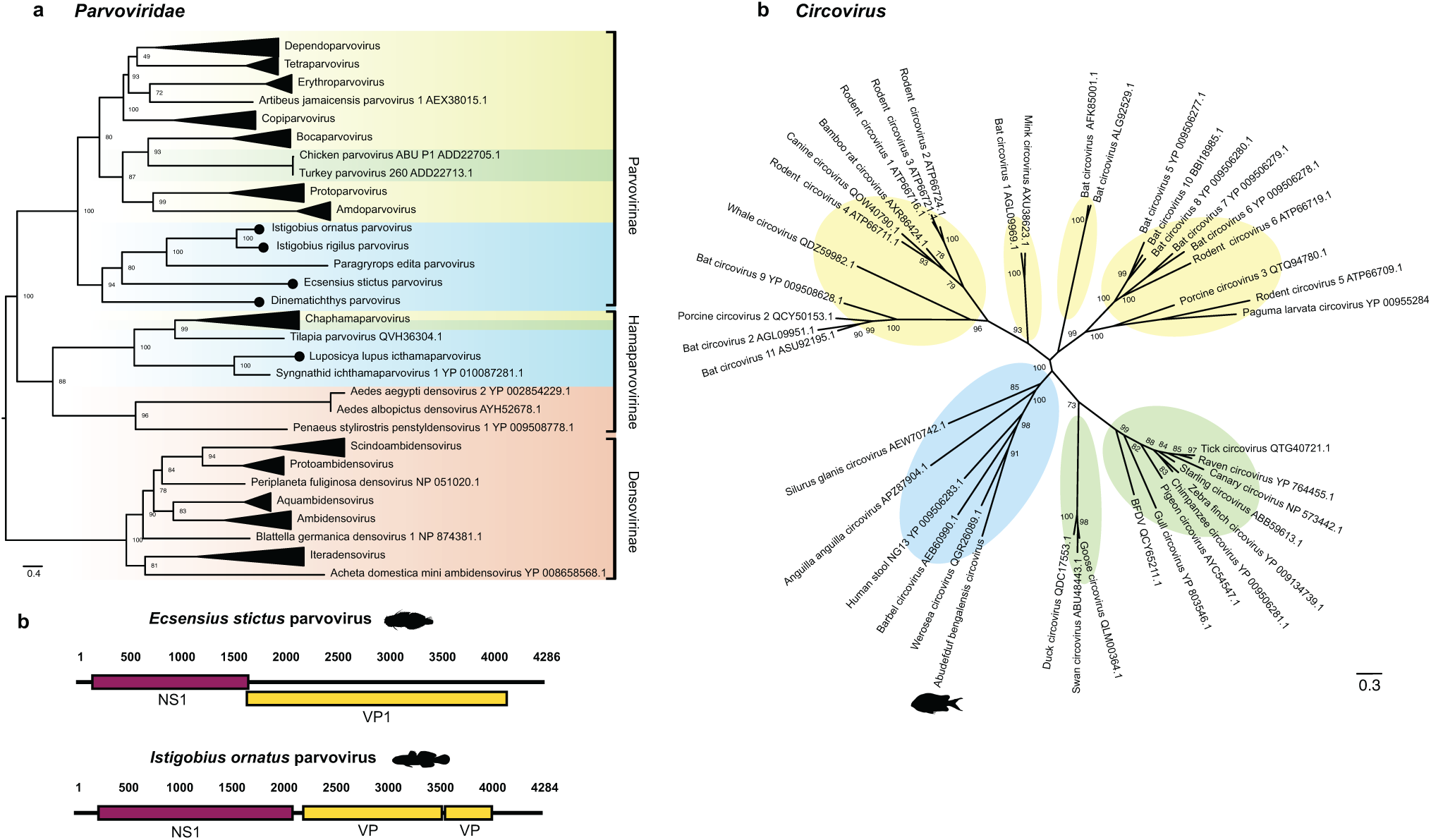
Phylogenetic relationships of the ssDNA viruses identified in this study (a) Phylogenetic analysis of the NS1 gene of the *Parvoviridae*. Discovered viruses are represented as black circles. The tree is midpoint rooted for clarity. Scale bar represents the number of amino acid substitutions per site. (b) Schematic genomes (nt) of discovered parvoviruses. Coloured boxes represent both structural (NS1) and non-structural (VP) ORFs. (c) Unrooted phylogeny of the replication-associated protein of the genus *Circovirus* (*Circoviridae*). Discovered circovirus is represented as a fish symbol. Branches of both trees are highlighted to represent host class: blue, fish; red, invertebrates; green, birds; yellow, mammals.

We also identified the partial genome of a novel icthamaparvovirus in *L. lupus* and a novel circovirus in *Abudefduf bengalensis* that clustered within a distinct clade of fish circoviruses. (Figure 4).

### dsDNA viruses

Analysis of the *Halichoeres melanurus* meta-transcriptome identified nine conserved proteins sharing 99-100% similarity with all currently described *Santee-Cooper ranavirus* isolates such as largemouth bass virus, manadarin fish ranavirus, koi ranavirus, doctor fish virus, and guppy virus 6 (*Iridoviridae*; Supplementary Table 2). We used the highly conserved major capsid protein and DNA polymerase for phylogenetic analysis [50], which further confirmed a novel *Santee-cooper ranavirus* isolate, tentatively named *H. melanurus* ranavirus (Figure 5). Largemouth bass virus and mandarin fish ranavirus are highly lethal in farmed populations, causing 95-100% mortality [60, 61]. Our metatranscriptomic analysis also identified a novel iridovirus in *E. tutuilae* that fell sister to erythrocytic necrosis virus (ENV) [51] and clustered with other erythrocytic-like viruses identified in amphibians and reptiles [52, 53, 54, 55]. ENV was the closest relative across all proteins identified (Supplementary Table 2).

**Figure 5.**
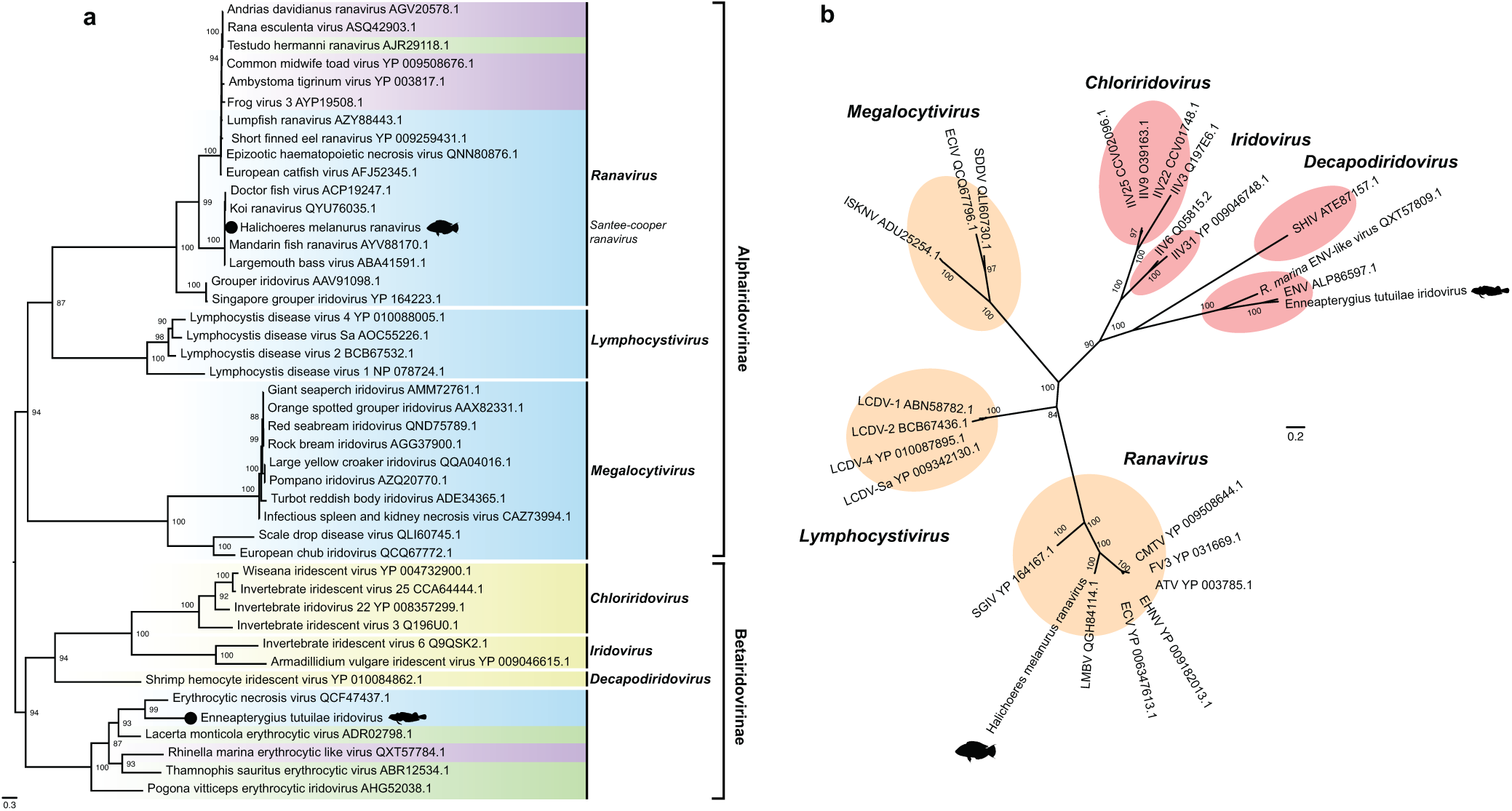
(a) Phylogenetic analysis of the DNA polymerase gene among the *Iridoviridae*. Scale bar represents the number of amino acid substitutions per site. Tip labels represent virus name with NCBI/GenBank Accession. Viruses discovered here are represented as a black circle and fish symbol. Branches are highlighted to illustrate host class: blue, fish; purple, amphibians; green, reptiles; yellow, invertebrates. (b) Unrooted phylogeny of the MCP of the *Iridoviridae*. Branches are coloured to represent both subfamilies: orange, *Alphairidovirinae*; red, *Betairidovirinae*.

### Cross-species virus transmission in a reef fish community

Despite the large number of viruses identified, we only found evidence for one cross-species transmission within our GBR ecosystem. This involved astroviruses found in five different fish species that exhibited high levels of amino acid sequence similarity and phylogenetic clustering (Figure 3). Specifically, phylogenetic comparisons of ORF1b (RdRp) revealed two viral species: goby astrovirus 1, identified in *I. goldmanni*, and goby astrovirus 2, identified in *I. nigroocellatus, I. decoratus, Asterropteryx semipunctatus*, and *Cabillus tongarevae* (Figure 4). These two viruses exhibit 82.5% amino acid sequence similarity across the virus genome, while the four sequences of goby astrovirus 2 had 96.8% amino acid similarity (Figure 3).

We also detected related viruses (i.e., those from the same virus family) in several different fish species, including the *Hantaviridae, Rhabdoviridae, Paramyxoviridae*, and *Picornaviridae* (Figure 6). However, most of these were highly divergent and likely reflect common ancestry rather than direct host-jumping in the reef ecosystem. For example, out of the eight picornaviruses identified in this study, the closest relatives were *Asterropteryx spinosa* picornavirus and *A. semipunctatus* picornavirus that shared only 44% amino acid similarity.

**Figure 6.**
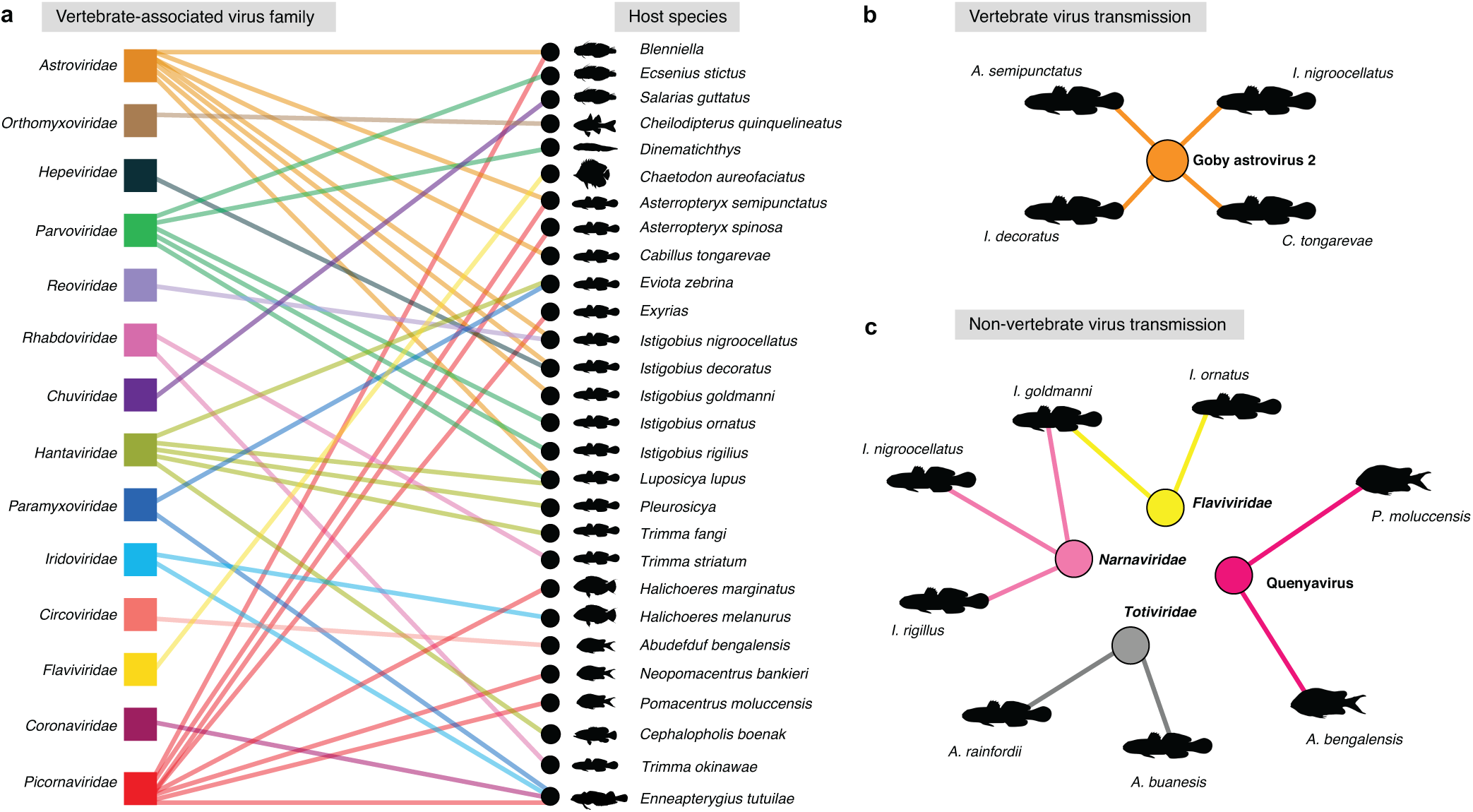
(a) Bipartite network illustrating viral families shared between fish taxa. Viral families are represented as a coloured box, with viruses connected between host species as a coloured line. (b) Network diagram illustrating vertebrate-associated viral species shared between fish. Coloured circle and lines illustrate viral species. (c) Network diagram revealing non-fish viral species shared between fish libraries. All fish silhouettes broadly represent host species at the family level.

Also of note was the identification of viral species not directly infecting the fish themselves, but rather associated with the local environment, diet or microbiome (i.e., non-fish) that were transmitted between reef fish assemblages. These were quenyaviruses (95% amino acid similarity between *Pomacentrus moluccensis* and *A. bengalensis*), flavi-like viruses (91.2% between *I. goldmanni* and *I. ornatus*), narnaviruses (97.7% between *I. goldmanni, I. nigroocellatus* and *I. rigillus*) and totiviruses (93.5% between *Amblygobius buanesis* and *Amblygobius rainfordi*) (Supplementary figs 1, 2 and 4; Figure 6). That there were more instances of cross-species transmission of non-fish viruses compared to those viruses that actively replicate in fish suggests that the latter group are subject to strong host barriers, even among closely related species.

### Comparisons of viral alpha and beta diversity between reef fish families

We next compared vertebrate virome composition between reef fish families, as well as between cryptobenthic reef fishes and large reef fishes (i.e. that differ in size). In our data set, cryptobenthic reef fish families included the Gobiidae, Apogonidae, Blenniidae, Tripterygiidae, Bythitidae and Pseudochromidae, while the large reef fish families included the Pomacentridae, Acanthuridae, Tetraodontidae, Atherinidae, Serranidae, Monacanthidae, Chaetodontidae, Labridae, Muraenidae, and Ophichthidae [10].

Three statistical measures were used to assess alpha diversity: viral abundance (i.e., standardised number of viral reads), observed viral richness (i.e., the number of viruses) and Shannon diversity. Notably, we found an association between fish size and observed viral richness, with cryptobenthic reef fishes harbouring more viruses than large reef fishes (χ^2^ = 2.795, df = 1, P = 0.028). In particular, the Tripterygiidae exhibited greater observed viral richness than all other reef fish families (χ^2^ = 16.678, df = 15, P = 0.007). However, we found no association between reef fish family and viral abundance (P = 0.153), Shannon diversity (p = 0.901) or beta diversity (R^2^ = 0.334, P = 0.064). Likewise, we identified no significant differences in viral abundance (P = 0.271), Shannon diversity (P = 0.142) nor beta diversity (R^2^ = 0.048, P = 0.121) between cryptobenthic reef fishes and large reef fishes.

As a form of internal control, we repeated our analyses of viral abundance and diversity on the non-fish viruses identified here. This analysis revealed no significant differences in viral abundance between reef fish families (P = 0.994) nor between cryptobenthic reef fishes and larger reef fishes (P = 0.355). Similarly, we found no significant difference in observed viral richness between fish families (P = 0.733), although cryptobenthic reef fishes exhibited higher observed non-vertebrate viral richness than large reef fish families (χ^2^ = 10.805, df = 1, P = 0.016). We observed no difference in Shannon diversity between fish families (P = 0.453), as well as between cryptobenthic reef fishes and larger reef fishes (P = 0.070). Finally, we found no association between beta diversity and reef fish families (R^2^ = 0.279, P = 0.126) nor between cryptobenthic reef fishes and large reef fishes (R^2^ = 0.335, P = 0.058).

## Discussion

The GBR supports over 1200 species of fish and is the largest coral reef ecosystem in the world, comprising 2500 reefs across approximately 344,400 km^2^ [5]. Despite the global importance of the GBR, little is known about the natural diversity of viruses that infect tropical reef fishes, as well as the ecological and evolutionary processes that allow such viruses to spread within a reef fish community. To this end, we employed metatranscriptomic sequencing to characterize the viromes of 61 tropical reef fish species, including those occupying a 100 m^2^ reef fish community. This identified a total of 132 viruses, including 38 vertebrate-associated viruses and 94 that infected a broad range of invertebrate and fungal hosts.

Importantly, while we sampled coral reef fishes from a small spatial area, we identified a marked absence of cross-species virus transmission, with the only instance of host-jumping being the presence of a single viral species (*Astroviridae*) in four goby species (Figures 3 and 6). However, the variation across the four viral genomes is indicative of past host-switching throughout evolutionary history, which may span millions of years, rather than direct host-jumping within the ecosystem sampled here. Given the restricted spatial area considered, and the density of potential hosts, such a lack of cross-species transmission suggests that there may be important host genetic barriers to virus switching among the reef fishes sampled herein. This is supported by the observation that non-fish viruses – that are not impacted by aspects of host genetics – were characterized by higher levels of cross-species virus transmission. Although currently of unknown nature, these barriers are likely to be subtle and may reflect nuances in host cell receptor binding [56]. For instance, although the betacoronavirus RaTG13 sampled from *Rhinolophus affinis* bats is closely related (~96% sequence similarity) to SARS-CoV-2, it is unable to bind to the human ACE2 receptor [73].

Another notable result was that we detected differences in vertebrate virome composition between reef fish families, with cryptobenthic reef fishes harbouring more viruses than large reef fishes (Figure 7). Due to their small body size, cryptobenthic reef fishes exhibit significantly higher rates of metabolism compared to large reef fishes, resulting in high energy demands with a low tolerance for starvation [10]. Given their extremely short lifespans, a high degree of energy is used for their rapid growth as well as their complex reproductive strategies (e.g. sex change), such that there might be important energy trade-offs between growth, reproduction and immunity; particularly as immune systems require a substantial allocation of resources for recognizing and eliminating pathogens [10, 70]. While it is unclear if and how such energy demands impact immunity in cryptobenthic reef fishes, it is notable that this lifestyle is associated with a more diverse virome compared to large reef fishes, suggesting these species may be more susceptible to infection. Indeed, with a daily mortality rate of 8 to 9% [20, 76] investing in immunity may be a poor strategy, and it is possible that these energy-saving strategies are key to the spectacular success of these cryptobenthic fishes [77]. Whether as simple larvae in the pelagic realm, or on the reef, cryptobenthic fishes may be able to minimise energetically expensive activities such as immunity and swimming, freeing up energy to maintain adequate fecundity, despite exceptionally high mortality rates and infection risks.

**Figure 7.**
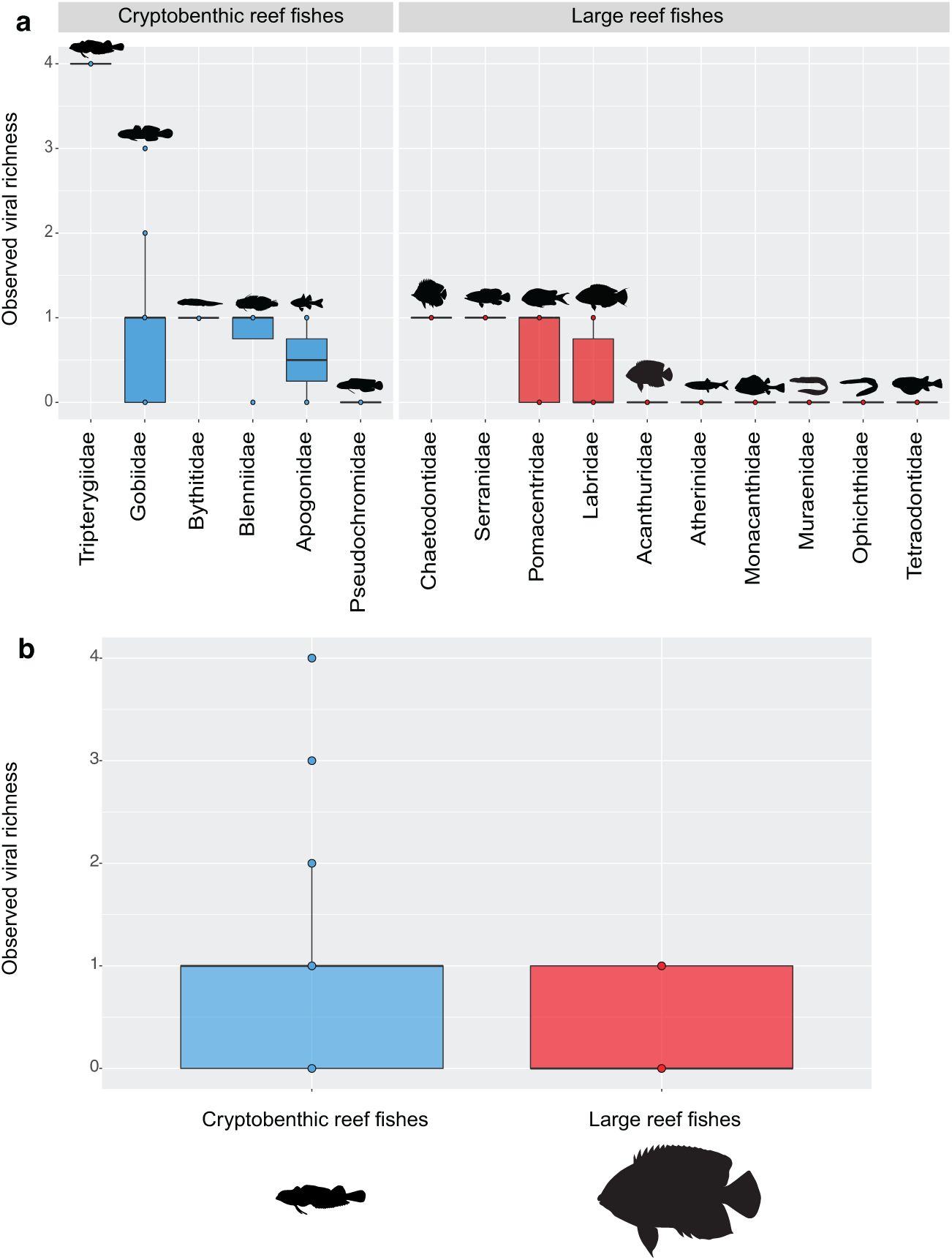
(a) Vertebrate-associated viral richness observed in all tropical reef fish families Comparison of observed vertebrate-associated viral richness between cryptobenthic reef fishes and large reef fishes.

Among reef fish families, the Tripterygiidae exhibited the greatest observed viral richness, with *E. tutuilae* harbouring four viral species. *E. tutuilae* is a generalist on coral reefs, utilising sand and rubble, soft coral, cave and open reef microhabitats [67]. It is therefore possible that habitat generalism may increase interactions with other fish species and hence increase the likelihood of being infected by a larger number of viruses. In contrast, *Gobiodon* species are extreme habitat specialists and may often only occupy a single coral species as habitat for its entire life with minimal interactions with other fish species [10, 68]. This lifestyle may explain why all five *Gobiodon* species examined here harboured no vertebrate-associated viruses, as well as few non-vertebrate associated viruses (Figure 1).

Given the phylogenetic distance between fish and other vertebrate classes, as well as the long-term associations through virus-host co-divergence [47, 57], it was not unexpected that 95% of the vertebrate-associated viruses discovered here clustered with other fish viruses. A case in point comes from the phylogeny of the *Hantaviridae* that clearly reflects the broad evolutionary history of vertebrates with the *Agantavirinae* (jawless fish) falling basal to the *Actantavirinae* (ray-finned fish), *Reptantavirinae* (reptiles) and *Mammantavirinae* (mammals), respectively (Figure 2). Despite their likely long-term presence in ray-finned fish, actinoviruses have only recently been discovered, and have been associated with disease in farmed species [19, 27, 44, 47]. For instance, a novel actinovirus – perch actinovirus – was identified in diseased European perch (*Perca fluviatilis*) with high concentrations of viral RNA in gill endothelial cells and macrophages [44]. Our discovery of four actinoviruses in gobies suggests these viruses may be widespread in this fish family and hence should be monitored closely if interacting with farmed populations.

Another virus of concern identified in our study is a novel isolate of *Santee-cooper ranavirus* in *H. melanurus*. To the best of our knowledge, this is the first discovery of this virus in Australia. Our detection of this virus in seemingly healthy wrasses as well as its natural presence in cleaner wrasses (*Labroides dimidatus*), suggests this family of tropical reef fishes may be important reservoir hosts for the *Santee-cooper ranavirus* group [63]. For instance, disease outbreaks with high mortality rates have only been observed in farmed freshwater fish species including largemouth bass (*Micropterus salmoides*), mandarinfish (*Siniperca chuatsi*) and koi (*Cyprinus carpio*) [58, 59, 60, 61, 62].

Despite the long-known presence of viral erythrocytic necrosis in various fish species across the North Atlantic and Pacific Oceans, the genome of the causative virus – ENV – was only recently characterized [51]. We identified a closely related virus in *E. tutuilae*. Interestingly, viral erythrocytic necrosis in a juvenile triggerfish (*Rhinecanthus aculeatus*) has been reported nearby at Lizard Island [64], suggesting that this virus may be circulating between several reef locations along the GBR. Phylogenetic comparisons of DNA polymerase and major capsid protein revealed a distinct ‘ectothermic vertebrate’ clade that could be clearly classified as a novel genus within the *Betairidovirinae, Iridoviridae* (Figure 5).

While we observed significant differences in virome composition between coral reef fishes, there were necessary limitations in our sampling. For instance, cryptobenthic fishes made up 62% of the fish diversity in this study. In addition, 23 libraries contained only one individual. Such unequal sample sizes likely impacted the statistical power of our analyses. Accordingly, future work should balance the number of samples as well as increase the number of reef sampling sites. Comparing virome composition between entire reef fish communities is crucial to understanding how host community diversity affects viral dynamics and disease emergence, particularly in the context of biodiversity loss. As such, larger ecological studies are needed to better evaluate whether a dilution or amplification effect occurs in a reef fish community.

In sum, our study identified a large diversity and abundance of viruses in tropical reef fish assemblages. Notably, we identified a marked absence of virus exchange within a reef fish community, suggesting there may be important host genetic barriers for successful cross-species virus transmission. Importantly, our discovery of a novel *Santee-cooper ranavirus* isolate in seemingly healthy wrasses, highlights the importance of virological surveillance in marine wildlife, particularly as this virus can cause significantly high mortality in farmed fishes. As such, these species should be considered in biosecurity risk assessments and screened if utilized for aquaculture or aquarium operations [69]. Accordingly, future studies should also investigate its susceptibility in important Australian food fish to fully assess its emergence threat [63]. Overall, this study increases our knowledge on the severely understudied coral reef fish virome and provides the first data on virus-host interactions in a reef fish community.

## Acknowledgements

This work was funded by Australian Research Council (ARC) Australian Laureate Fellowships to ECH (FL170100022) and DRB (FL190100062), and an ARC Discovery Project grant to ECH and JG (DP200102351). JLG is funded by a New Zealand Royal Society Rutherford Discovery Fellowship (RDF-20-UOO-007) and a Marsden Fast Start grant (20-UOO-105).

## Data availability

All sequence reads are available on the NCBI Sequence Read Archive (SRA) under BioProject XXXX (submission: SUB11462050, awaiting ID) and all generated virus genetic sequences have been deposited in NCBI/GenBank under accessions XXXX - XXXX (submission: 2582698, awaiting accession IDs).

## Supplementary information

**Supplementary Table 1. Host and meta-transcriptomic library information**

**Supplementary Table 2. Description of novel viruses identified in this study**

**Supplementary Figure 1**. (a) Phylogenies of the RdRp gene of +ssRNA non-vertebrate associated viruses in fish metatranscriptomes. Discovered viruses in this study are represented as black circles. Maximum likelihood trees were midpoint rooted for clarity only. Scale bar represents amino acid substitutions per site. Tip labels represent virus name with NCBI/GenBank accession.

**Supplementary Figure 2**. (a) Phylogenies of the RdRp gene of +ssRNA non-vertebrate associated viruses in fish metatranscriptomes. Discovered viruses in this study are represented as black circles. Maximum likelihood trees were midpoint rooted for clarity only. Scale bar represents amino acid substitutions per site. Tip labels represent virus name with NCBI/GenBank accession.

**Supplementary Figure 3**. (a) Phylogenies of the RdRp gene of −ssRNA non-vertebrate associated viruses in fish metatranscriptomes. Discovered viruses in this study are represented as black circles. Maximum likelihood trees were midpoint rooted for clarity only. Scale bar represents amino acid substitutions per site. Tip labels represent virus name with NCBI/GenBank accession.

**Supplementary Figure 4**. (a) Phylogenies of the RdRp gene of dsRNA non-vertebrate associated viruses in fish metatranscriptomes. Discovered viruses in this study are represented as black circles. Maximum likelihood trees were midpoint rooted for clarity only. Scale bar represents amino acid substitutions per site. Tip labels represent virus name with NCBI/GenBank accession.

**Supplementary Figure 5**. (a) Phylogenetic tree of the VP2 (RdRp) gene of the subfamily *Spinareovirinae* (*Reoviridae*). Discovered virus represented as a black circle. The tree is midpoint rooted for clarity only. Scale bar represents amino acid substitutions per site and tip labels display virus name with NCBI/GenBank accession. Branches are highlighted to illustrate host class: blue, fish; green, birds; red, mammals.

