## Supplementary figures and images for "Limited Cross-Species Virus Transmission in a Spatially Restricted Coral Reef Fish Community"

### Supplementary Figure 1

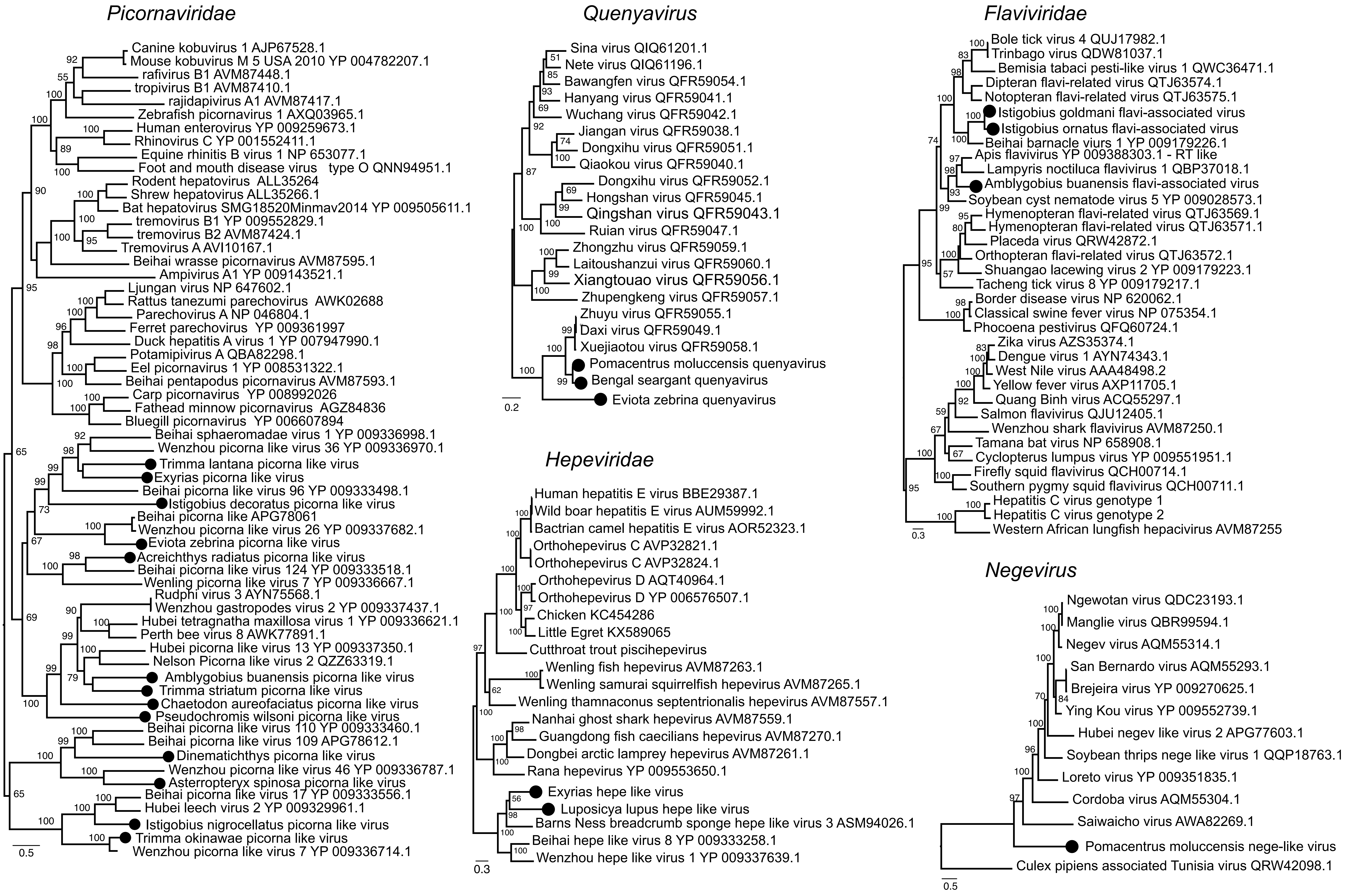

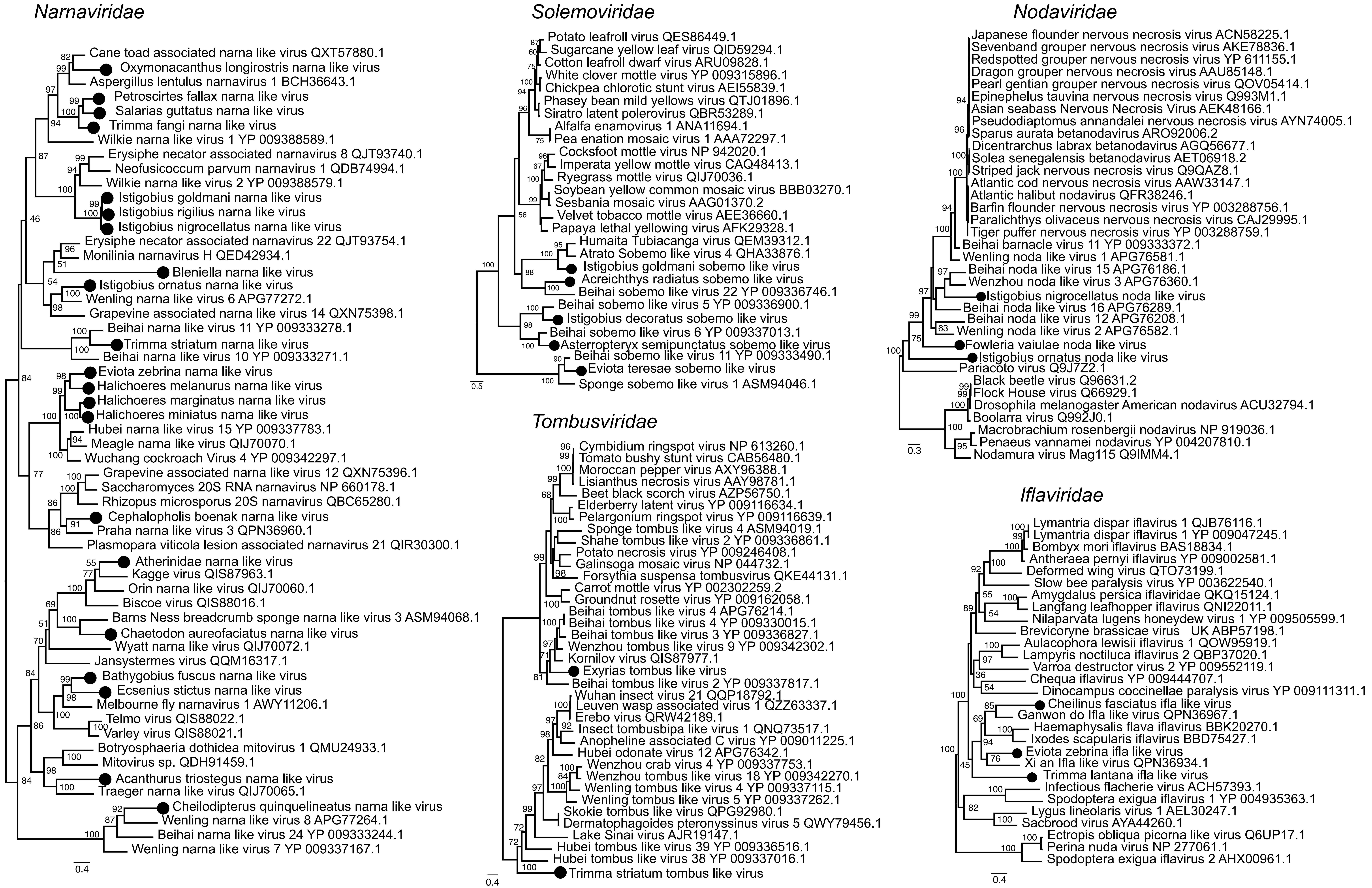

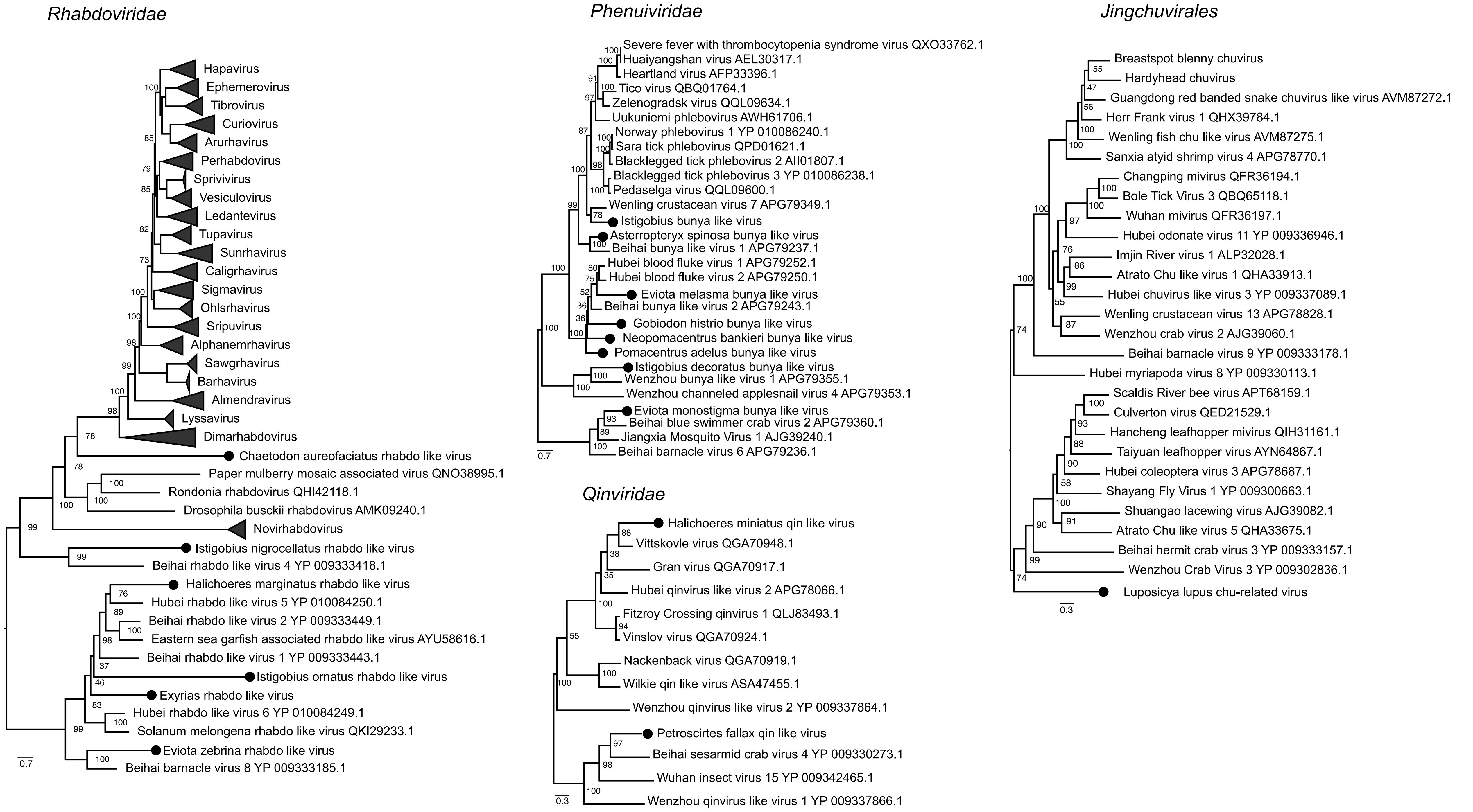

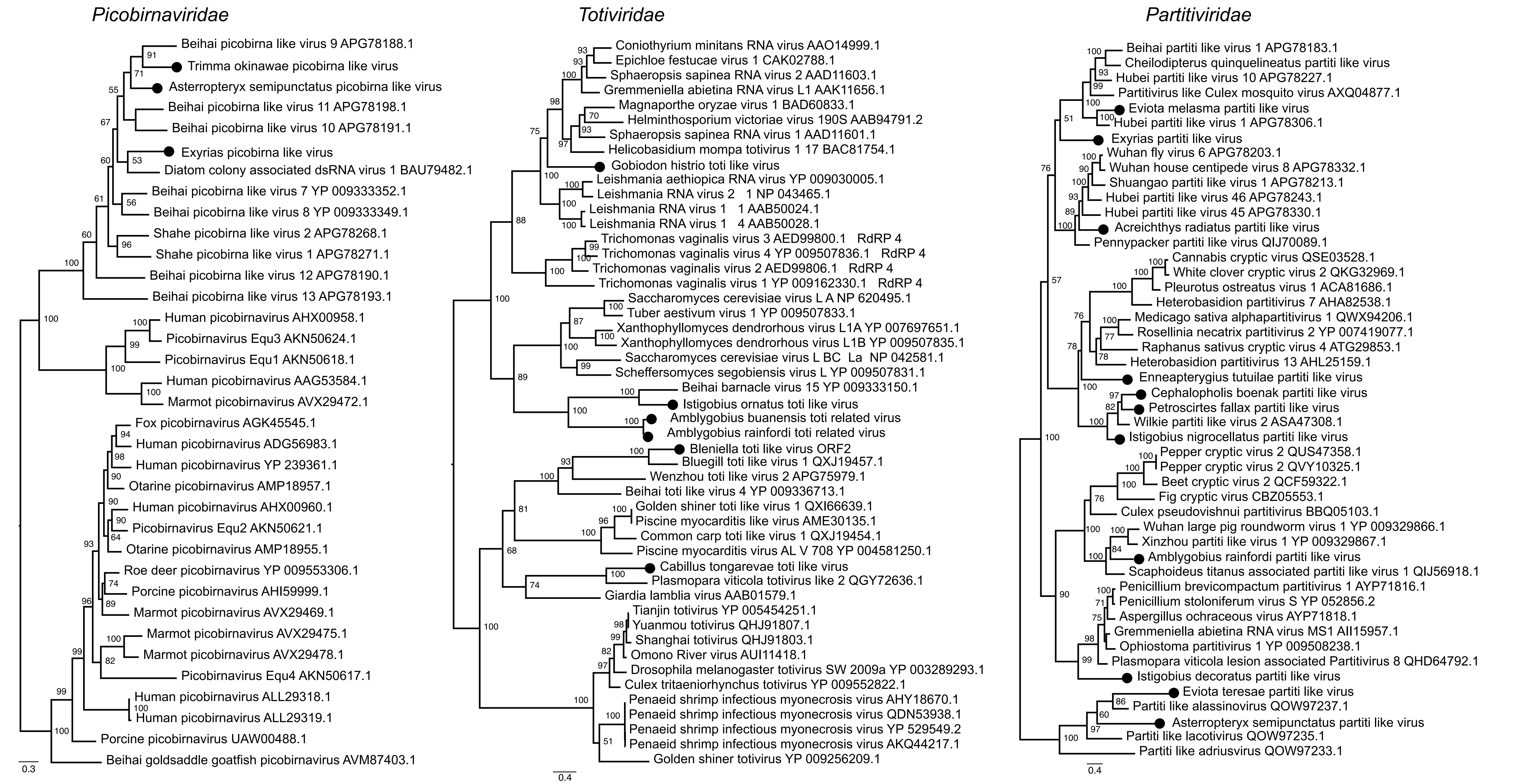

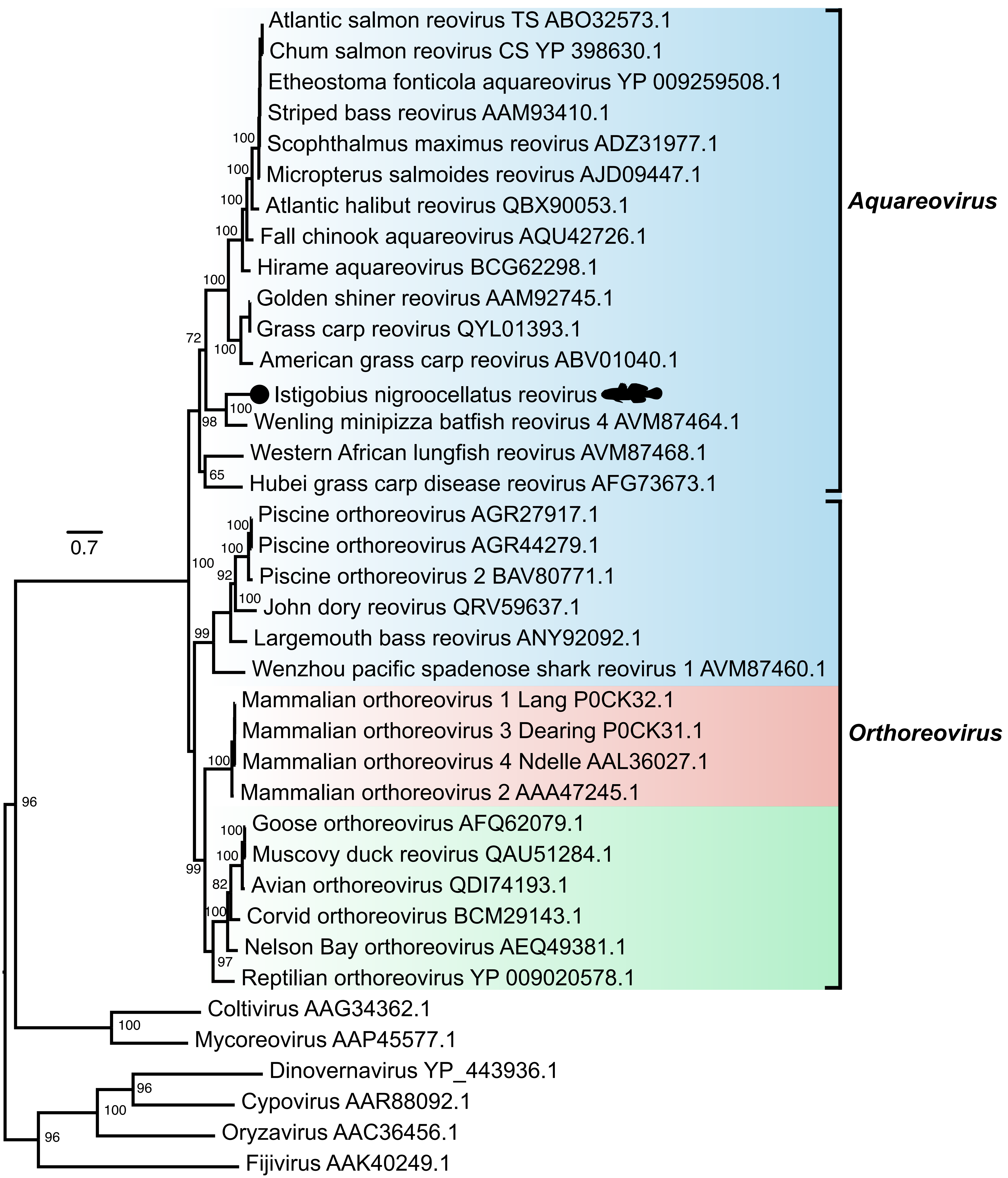

### Supplementary Figure 2

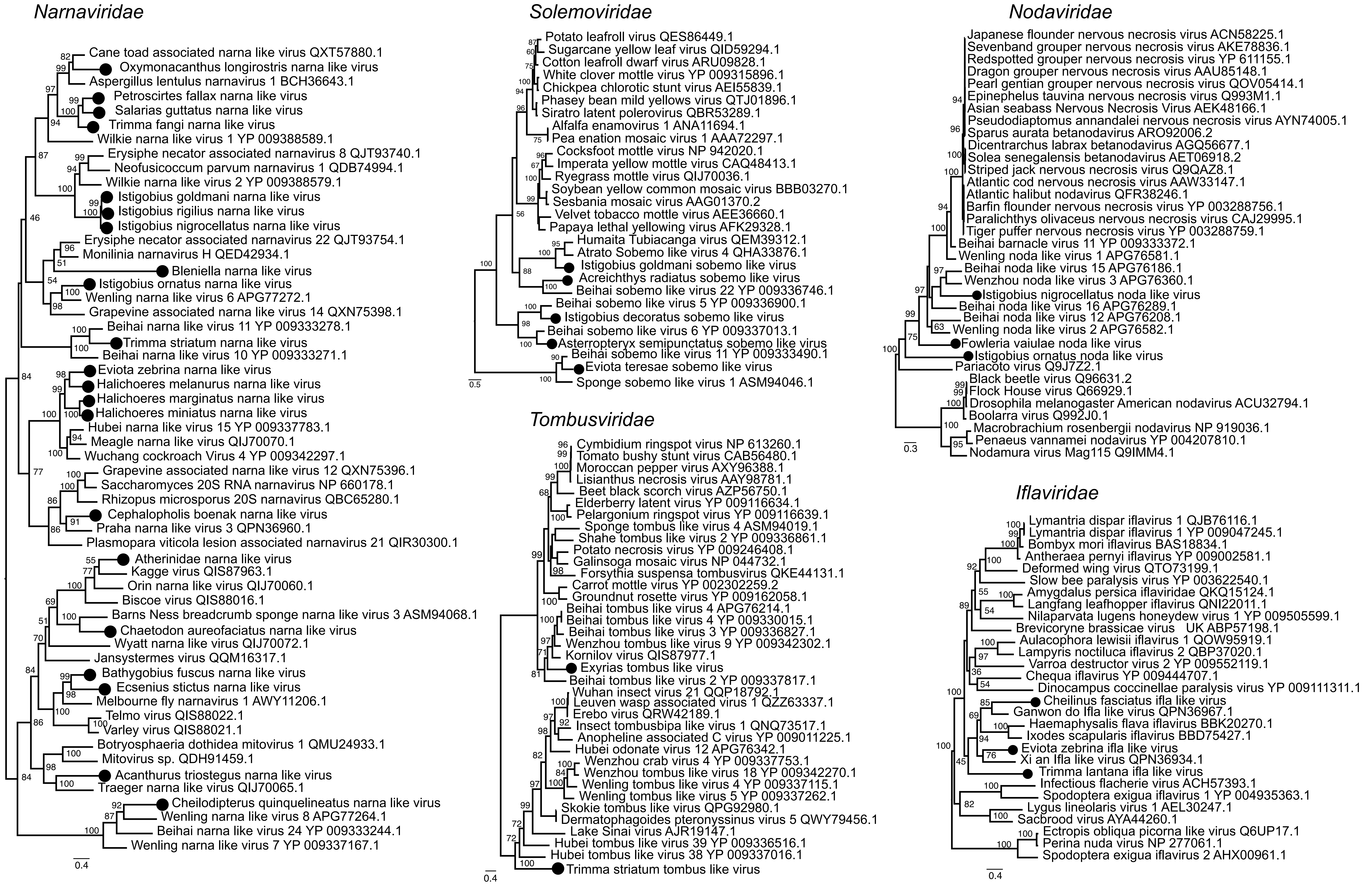

### Supplementary Figure 3

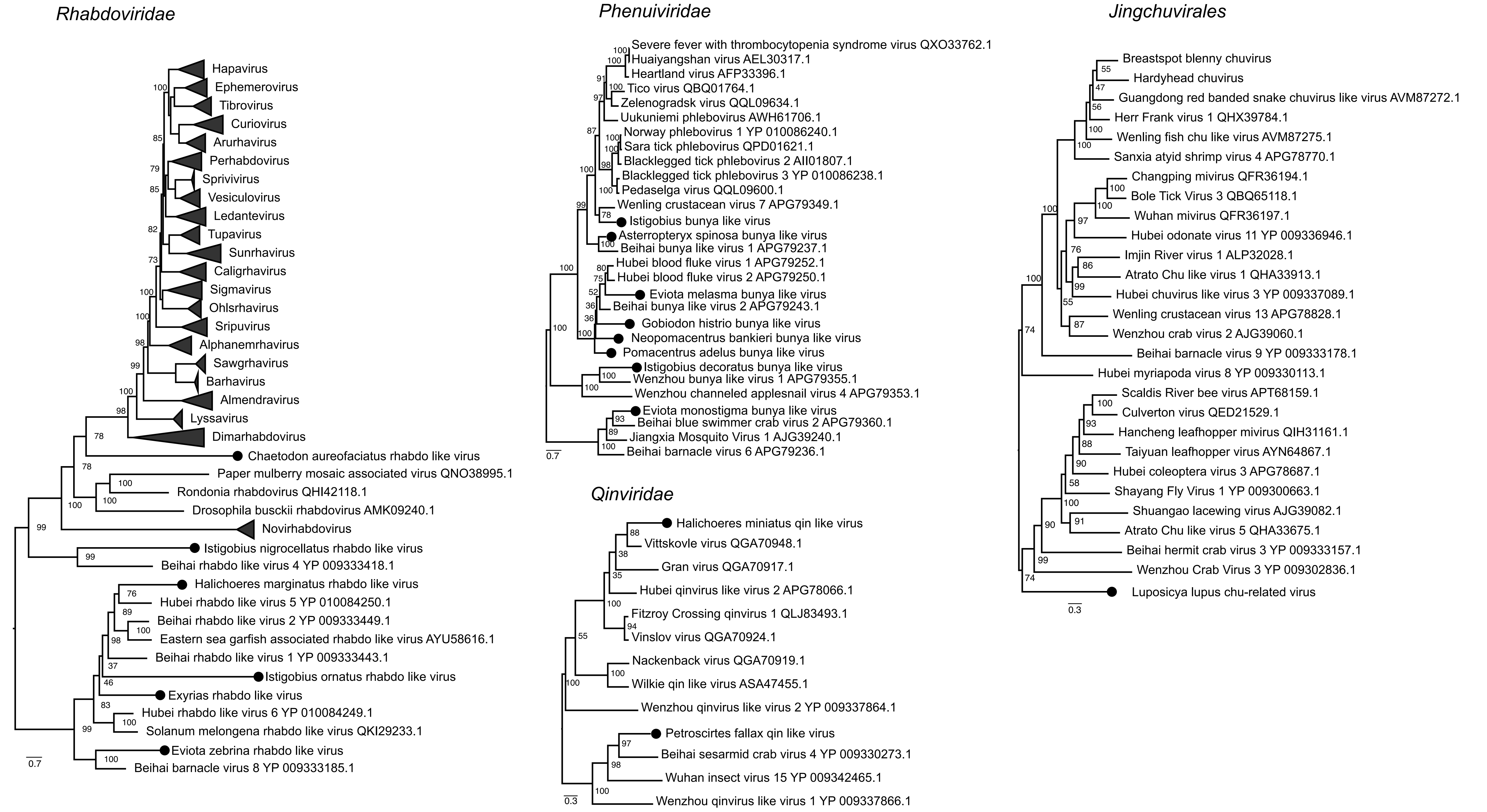

### Supplementary Figure 4

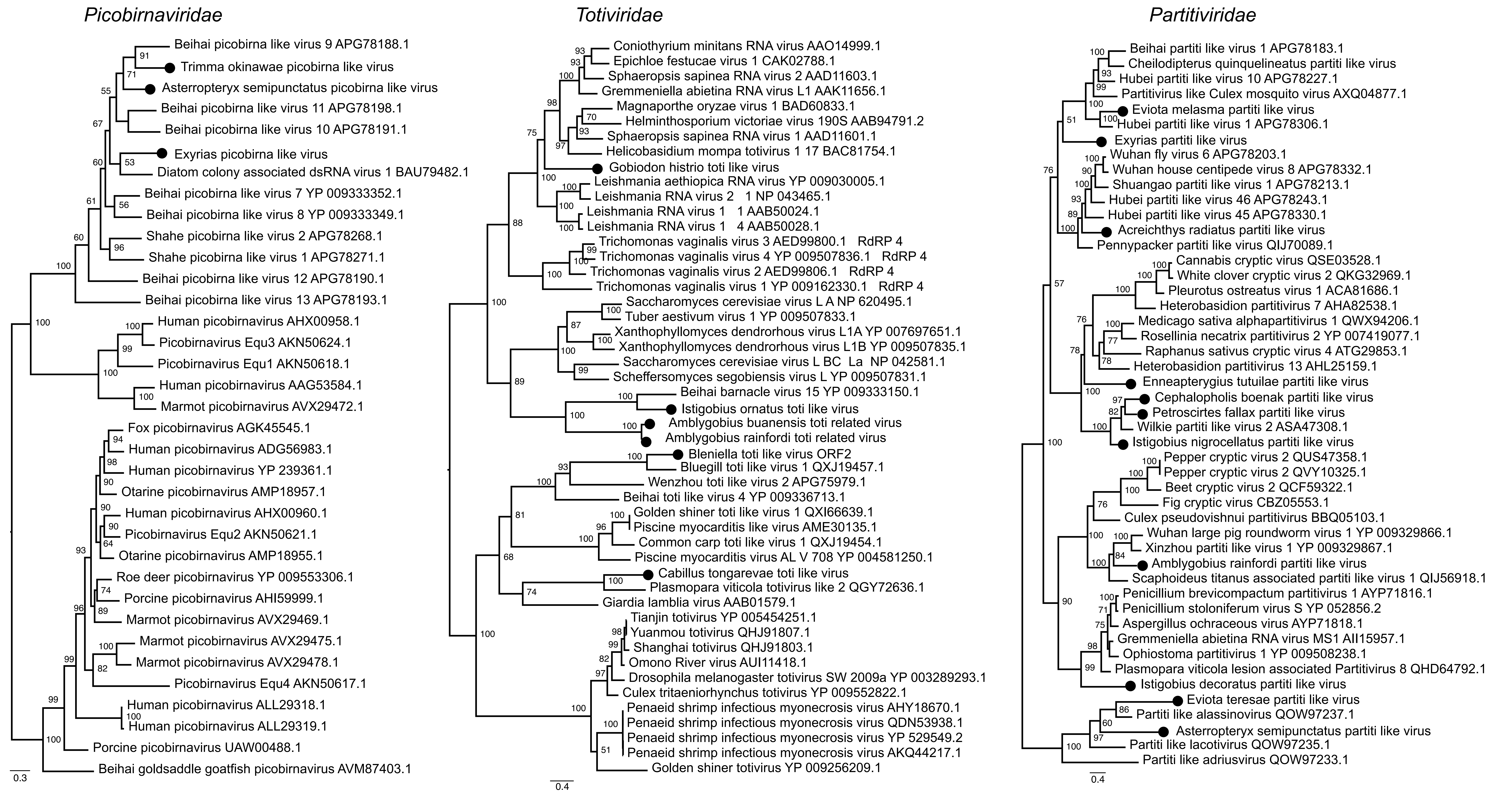

### Supplementary Figure 5

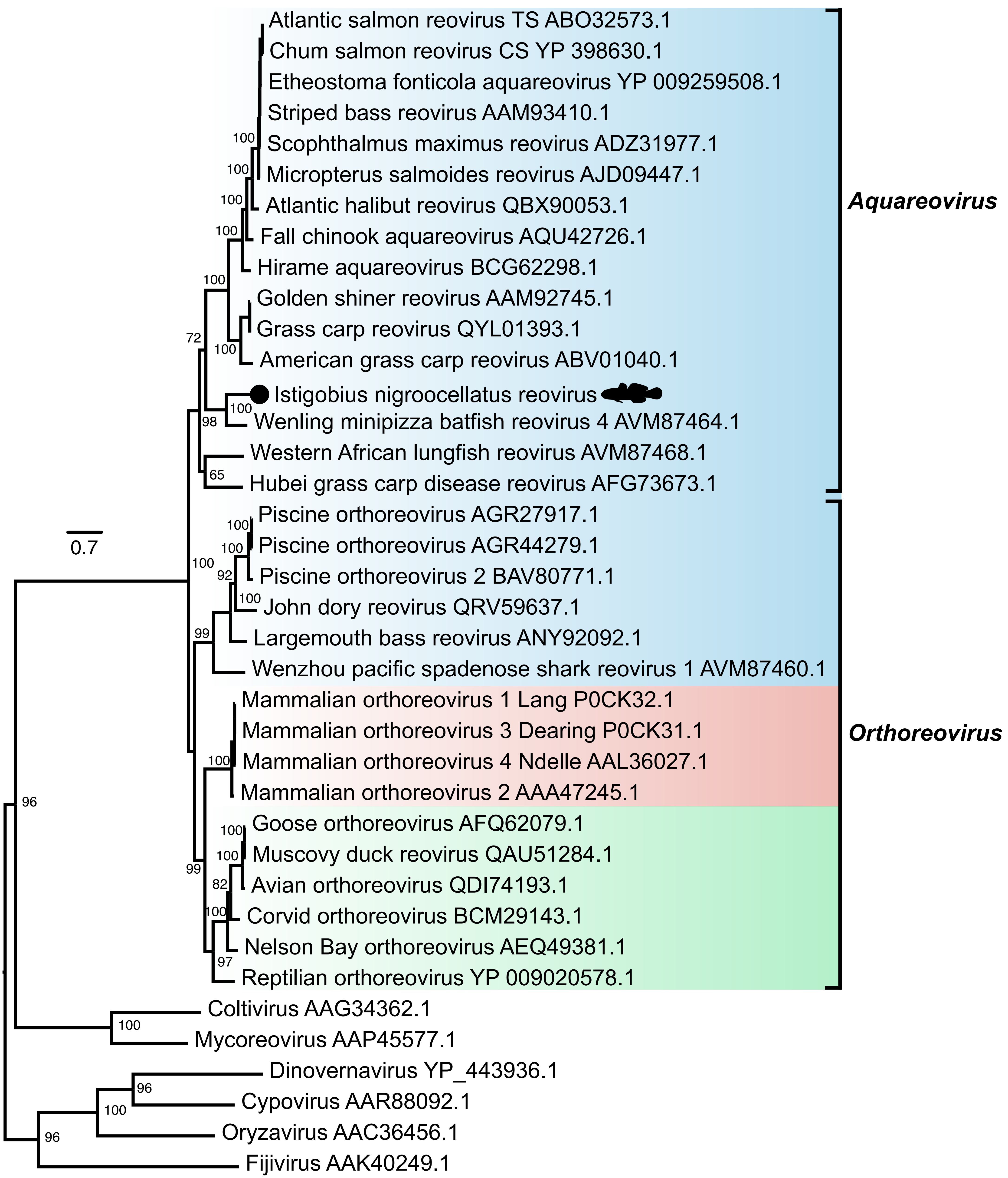
